# Vitrification by high pressure freezing of a wide variety of sample using the HPM Live µ

**DOI:** 10.1101/2024.03.06.583669

**Authors:** Chie Kodera, Yann Bret, Frederic Eyraud, Jérôme Heiligenstein, Martin Belle, Xavier Heiligenstein

**Affiliations:** CryoCapCell, Le Kremlin-Bicêtre, France

**Keywords:** Vitrification, high pressure freezing (HPF), electron microscopy (EM), transmission electron microscopy (TEM), freeze substitution, resin embedding

## Abstract

This study explores the efficacy and reliability of high-pressure freezing (HPF) as a sample preparation technique for electron microscopy (EM) analysis across a diverse range of biological samples. Utilizing the HPM Live µ technology, based on the historical hydraulic HPM010 from BalTec, we demonstrate the reliability of our industrial equipment to achieve the critical parameters necessary for vitrification. By directly measuring physical values within the HPF chamber, we ensure the proper functioning of the equipment, contributing to the technique’s reliability. A meticulous approach was adopted for each sample type, acknowledging the uniqueness of each specimen, and associating final sample analysis with its HPF curve, aiding in protocol optimization. Samples including human cell pellets, cell monolayer, mouse brain and liver biopsies, *C. elegans*, zebrafish, and *A. thaliana* root and seedlings were processed for EM analysis following HPF. The ultrastructure of each sample type was rigorously examined, revealing homogeneous preservation and minimal ice nucleation artifacts. Challenges such as plant leaf vitrification were addressed, highlighting the importance of methodological adaptation. Overall, our findings underscore the robustness and versatility of our HPM Live µ in preserving biological ultrastructure, offering valuable insights for researchers employing EM techniques in diverse biological studies.

## Introduction

Vitrification through High Pressure Freezing (HPF) represents a pivotal technique for immobilizing biological samples, facilitating their analysis via electron microscopy (EM). By rapidly freezing specimens under high pressure, HPF capitalizes on the water phase diagram to prevent ice crystal formation, instead inducing the formation of amorphous ice—a glass-like structure[1], [2], [3]. Achieving optimal vitrification conditions necessitates reaching 2076 bars, where water solidifies only at −22°C [4], followed by rapid temperature decrease below −100°C [5]at rates exceeding 2000K/s[6]. Under these conditions, water transitions into a metastable amorphous state [5], characterized by density comparable to its liquid form, thereby preserving the ultrastructure of surrounding biological material.

To attain these extreme physical conditions, we employ a liquid nitrogen flow pressurized at 2100 bars, released within a mere 10 milliseconds onto the sample. To safeguard delicate biological material during this process, the sample is positioned between two small metal carriers, typically composed of aluminum or gold-coated copper. These carriers transmit pressure to the hydrated sample while aiding heat extraction through their thermal conductive properties [6], [7].

Following vitrification, samples can be prepared for subsequent analysis using electron microscopy techniques, notably freeze substitution (FS) and plastic embedded electron microscopy or cryo-electron microscopy (cryo-EM). These techniques yield high-resolution images of the sample’s ultrastructure, vital for elucidating the function and behavior of biological molecules and complexes[8].

HPF/FS offers several advantages over conventional chemical fixation and room temperature (RT) dehydration methods. Firstly, it preserves samples in a state closely resembling their native condition, enhancing the accuracy of structural and functional analyses. Secondly, it mitigates distortions that may arise during dehydration and embedding, common pitfalls in RT chemical fixation processes, particularly beneficial when studying wide sample areas. Nonetheless, HPF is acknowledged as a complex and expert technique, necessitating substantial optimization.

This article showcases the versatility of our newly developed high-pressure freezing machine, the HPM Live µ [9], across a diverse array of biological specimens commonly studied in research laboratories. Addressing a longstanding community need, our study provides a comprehensive and rigorously tested collection of protocols tailored to a modern HPF platform, with the aim of democratizing this technique.

All samples in our study were vitrified on the HPM Live µ using preset parameters. Standard aluminum carriers were employed, with details provided in Table 1. Cryoprotectants were utilized to minimize variability from the culture medium[10], [11], [12], as outlined in Table 1. Freeze substitution was conducted using standard cocktails, while embedding was performed in EPON resin, widely adopted in the field [13], [14], [15], [16]. Unstained sections of 70nm were subsequently imaged by TEM.

**Table 1.**
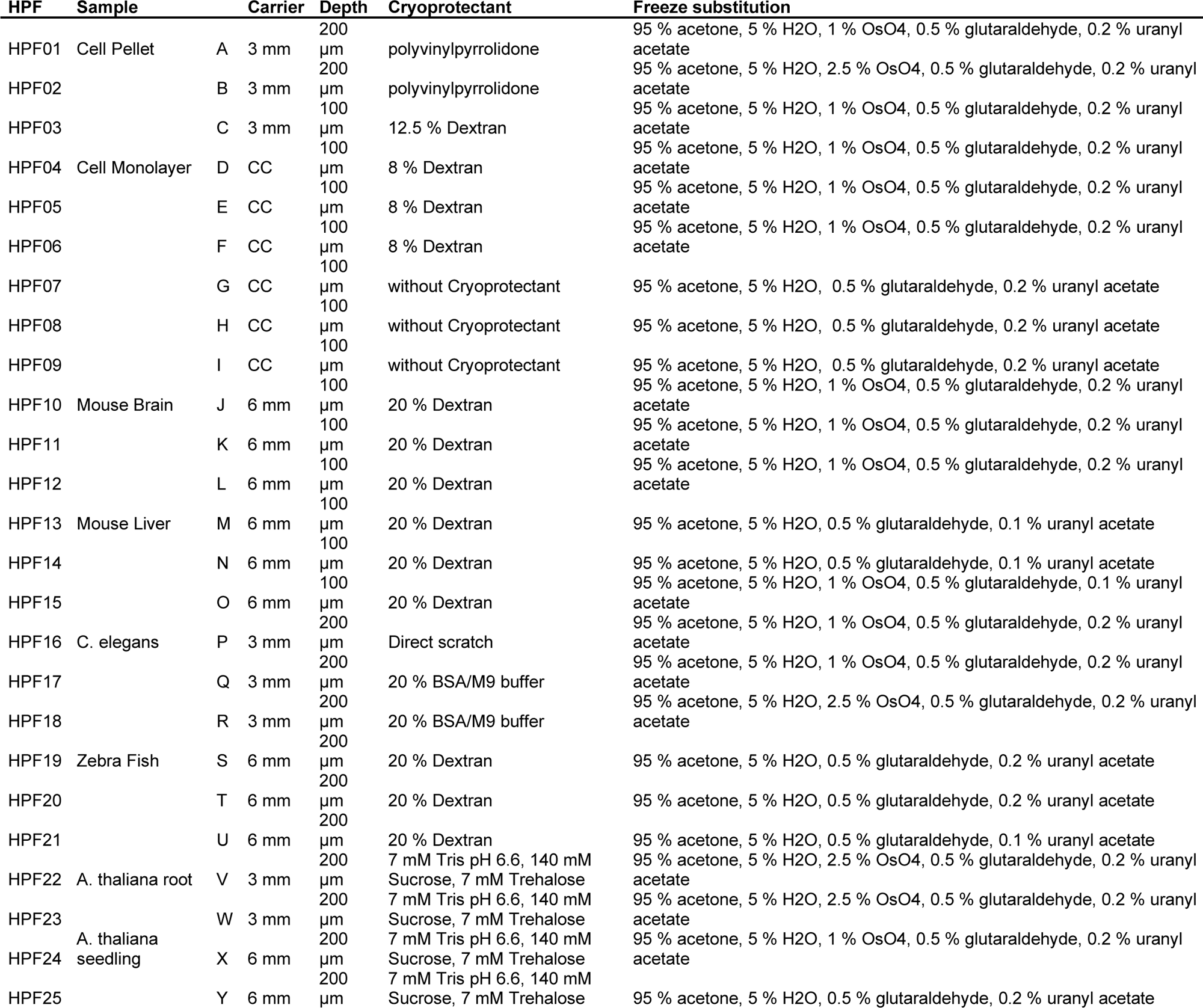
HPF conditions. HPF and freeze substitution conditions for 25 independent experiments.

We present this manuscript as a foundational reference for biological exploration, allowing scientists to utilize our protocol with confidence, knowing it can yield satisfactory preliminary results. Our images may serve as a reference point for project-specific optimization, advancing the collective understanding and application of HPF techniques in biological research.

## Materials and methods

### Sample preparation before High-Pressure Freezing

#### Generic care

Loading carriers with samples must be done with care. ***<u>No air bubble</u>*** should be left in the sample to avoid pressure plateau reduction during vitrification.

All carrier for experiment were prepared in advance to keep focus on the sample. No carrier coating was applied to limit variability among experiments.

B carriers, generally used to close the chamber, are placed flat side down, onto a filter paper impregnated with the appropriate cryoprotectant. This prewetting step reduces the formation of air bubble during cavity closure.

All samples were vitrified as fresh as possible to limit degradation prior to HPF.

All samples were prepared and vitrified in the shortest possible time, from their culture media to the aluminium carrier.

#### Monolayer Cell into a Pellet

Neuroblastoma cells, type SHSY-5Y, were cultured in complete medium (1x DMEM+GlutaMAX, 20 % fetal bovine serum, G418) at 37 degrees up to 70% confluence. Treated by Trypsin, resuspended in complete medium and pelleted by centrifugation. The cell pellets were resuspended in cryoprotectant (table 1) and then pipetted in an A carrier (3 mm) of 100 µm or 200 µm depth (table 1). The carrier was closed by a B carrier flat side to create a 100 µm or 200 µm deep cavity prior to loading into the HPM clamp for vitrification.

#### Monolayer Cell cultured onto a CryoCapsule

Neuroblastoma cells, type SHSY-5Y, were cultured in complete medium (1x DMEM+GlutaMAX, 20 % fetal bovine serum, G418) at 37 degrees up to 40% confluence into a CryoCapsule[17]. On the day of vitrification, the CryoCapsule was closed using a covering sapphire disc, loaded into the specific CryoCapsule clamp [18]. The cells involved in a correlative light and electron microscopy (CLEM) project were imaged at 5 or 20x prior to vitrification in absence of cryo-protectant (figure 4). Cells not involved in a CLEM project were exposed to 8% dextran, loaded as previously described and vitrified.

#### Mouse biopsy (Brain, and Liver)

F8-deficient mice obtained from Kazazian et al[19] were backcrossed (>10 times) on a C57Bl/6 background. Housing and experiments were performed following French regulations and the experimental guidelines of the European Community. Mice were deeply anaesthetized and subjected to intracardiac perfusion with 4 % paraformaldehyde in 150mM phosphate buffer prior to biopsy to ensure proper fixation of the tissues. Prefixation offers several advantages: it slows down sample degradation and enhances tissue rigidity, facilitating subsequent slicing steps using an oscillating knife (vibratome VTS 2000, Leica Microsystems). Tissues or organs were maintained at 4°C between biopsy and thinning. All manipulations, from thinning to vitrification, were conducted at room temperature.

Large organs were reduced to blocks no more than 5 mm in height and affixed onto the vibratome tray from their larger face using cyanoacrylate adhesive (super glue). Excess adhesive was carefully removed using filter paper, and the sample was submerged in PBS. It’s notable that super glue exhibits strong reactivity to PBS, and any excess may result in the generation of contaminating fibers in the bath, although this observation is not directly presented in our data. The amplitude and frequency of the razor blade in the vibratome were adjusted as needed until a smooth, gently floating 100µm slice of tissue was obtained. The floating slice was carefully moved to a clean area of the PBS bath using a fine paintbrush. A biopsy puncher was then used to extract a 2 or 4 mm disc from the slice, which was lifted out of the PBS bath with the paintbrush and deposited into the A carrier. A drop of cryoprotectant was added, and excess solution was removed with filter paper until a tiny concave meniscus remained. Subsequently, the sample was encapsulated by a flat side of the B carrier, which had been pre-wetted with the cryoprotectant solution to minimize air bubble encapsulation. The assembly was then loaded into the HPM clamp and subjected to vitrification. A short video is available on our website: https://www.cryocapcell.com/tips-and-tricks-for-em and on YouTube https://www.youtube.com/shorts/xz17Wy_l_64

### Caenorhabditis elegans

*C. elegans* (FL378 ERM-1 erm-1(bab59[erm-1::mNG^3xFlag]) I) [20] was maintained under typical conditions as described[21]. All manipulations were conducted at room temperature. Two strategies were employed to concentrate the sample in the carrier:

1. **Direct Scratch Method**: *C. elegans* were collected and concentrated by directly scratching the surface of the agar plate using a wooden toothpick, without the addition of cryoprotectant. The collected worms were then transferred to the carrier prior to sealing and vitrification. This method closely resembles the approach used for vitrifying yeast colonies from agar plates.
2. **Cryoprotectant Method**: 100µL of cryoprotectant solution consisting of 20% BSA in M9 buffer was prepared in a 1.5 ml Eppendorf tube. *C. elegans* were collected by gently scratching the surface of the culture plate with the open edge of the tube. The collected worms were concentrated via brief centrifugation. Subsequently, the mixture of *C. elegans* and cryoprotectant was aspirated using a micropipette and dispensed onto an A carrier (refer to Table 1). The carrier was then sealed with a flat side of the B carrier and subjected to vitrification.

These two approaches enabled efficient concentration of *C. elegans* samples within the carriers, facilitating sample retrieval in the electron microscope after random ultramicrotomy.

#### Zebrafish - *Danio rerio*-

Three-day-old (72 hour post fertilization) zebrafish *Tg (HuC:eGFP)* larvae reared under typical laboratory conditions were utilized for this study. Embryos were staged and cared for according to standard protocols (https://zfin.org/zf_info/zfbook/cont.html). All animal experiments were conducted with approved protocols at Inserm by DDPP Val de Marne, France, under license number F 94-043-013. All manipulations were conducted at room temperature. The zebrafish larvae were first anaesthetized in a 35 mm petri dish containing 3 ml of osmosis water supplemented with 10 drops of Tricaine 0.4%. After a 2-minute anaesthetization period, the larva was carefully transferred from the petri dish using a Pasteur pipette and placed onto an A carrier (refer to Table 1). Excess water was removed from the carrier using a filter paper, following which a drop of cryoprotectant (20% Dextran in osmosis water) was added. The carrier was then sealed with a flat side of the B carrier, which had been pre-wetted with the cryoprotectant solution, and subjected to vitrification.

#### *Arabidopsis thaliana* (seedlings and roots)

Three to five-day-old *A. thaliana* (*Col-0*) seedlings cultured on Murashige and Skoog medium [22]plates under room temperature with continuous light were utilized in this study. All manipulations were conducted at room temperature. The seedlings were gently removed from the culture plate using tweezers and carefully deposited onto a clean glass slide. To prevent desiccation, several drops of cryoprotectant solution (7 mM Tris pH 6.6, 140 mM Sucrose, 7 mM Trehalose) were added to cover the plants.

Subsequently, the seedling, or the root tip excised using a sharp flesh razor blade was transferred to an A carrier (refer to Table 1). Another drop of cryoprotectant was added to the carrier to ensure proper protection of the sample. The carrier was then sealed using a flat side of the B carrier, which had been pre-wetted with cryoprotectant, and subjected to vitrification.

### High-Pressure Freezing

Following the aforementioned preparations, each sample was subjected to vitrification using our HPM Live µ® apparatus (CryoCapCell, Paris, France). Prior to the initiation of the high-pressure liquid nitrogen flow, a brief pre-injection of ethanol lasting 75 (sample J, K, L) or 150 milliseconds (all other samples) was administered. This pre-injection serves to delay cooling while the system reaches the desired pressure. Throughout the process, temperature and pressure were directly measured inside the HPF chamber using sensors positioned above and below the specimen. Notably, no data manipulation or calculation was applied to the collected data, which is presented in its raw form within this manuscript. This unprocessed data serves to illustrate the repeatability of our apparatus and the precise trajectory of each sample across the water phase diagram.

Consistency and standardization were upheld by systematically applying the same HPF parameters to all samples, thereby minimizing variability in our protocol. This approach ensures robust and reproducible results across the experimental cohort.

### Freeze Substitution and embedding

Following vitrification, the samples underwent freeze substitution using an FS-8500 RMC (Boeckeler Instruments, Tucson, Arizona) in a solution mix detailed in Table 1. The freeze substitution process followed a temperature ramp program as outlined below:

For the standard freeze substitution protocol:

- 18 hours at −90 degrees
- 15 hours from −90 degrees to −60 degrees (with a temperature increase of 2°C per hour)
- 8 hours at −60 degrees
- 15 hours from −60 degrees to −30 degrees (with a temperature increase of 2°C per hour)

Upon completion of the program, the samples were kept on ice for 1 hour, followed by three washes in acetone. Subsequently, the samples were embedded in EPON 812 resin through a series of steps: 2 hours each at 30%, 60%, and 100% resin concentration, repeated twice. The embedding process concluded with 48 hours of polymerization at 45°C.

For the sample Q (*C. elegans*) variant (Table 1), the temperature ramp program differed as follows:

- 14 hours from −90 degrees to −40 degrees (with a temperature increase of 3.6°C per hour)

Following the completion of the temperature ramp program, the sample Q underwent the same post-substitution processing steps as described for the standard freeze substitution protocol.

### Ultra microtome

70 nm to 80 nm thin sections were made on a Ultracut S (Leica) with an oscillating diamond knife Ultra Sonic (Diatome, Switzerland) then collected on homemade formvar coated mesh grids.

### Transmission Electron Microscope Imaging

TEM imaging was performed with a Tecnai 12 transmission electron microscope, 120 kV (FEI) equipped with a CCD camera (OneView 4Kx4K Gatan) controlled by GMS software.

## Results

### Twenty-five independent HPFs with optimised conditions

High pressure freezing (HPF) is often perceived as a sophisticated sample preparation technique, subject to variability influenced by factors such as sample type, equipment, and operator proficiency. However, we assert that HPF is inherently reliable, with the initial mastery of sample preparation being the pivotal aspect achievable within a few trials. Leveraging our HPM technology, derived from the esteemed hydraulic HPM010 by BalTec, we consistently applied critical parameters to achieve vitrification. Furthermore, our direct measurement of physical values inside the HPF chamber serves as a valuable tool for assessing equipment functionality.

Recognizing the labor-intensive nature of obtaining electron microscopy (EM) images following vitrification, we refrained from pooling samples randomly and instead treated each sample as unique. Associating TEM analysis with its corresponding HPF curve proves invaluable when establishing protocols for novel sample types. A good HPF curve juxtaposed with a suboptimal image informs optimization efforts in sample preparation.

To validate this reliability, we rigorously tested a diverse array of samples, all prepared on the same apparatus by the same team of experimentalists. Our sample selection encompassed human cell cultures, mouse biopsies (including brain, and liver tissues), *C. elegans*, zebrafish, and *A. thaliana* (both root and seedling specimens), representing key inquiries in contemporary biology. Our comprehensive procedure is delineated in Figure 1, with vitrified samples undergoing freeze substitution, resin embedding in EPON, thin sectioning, and subsequent TEM observation.

**Figure 1.**
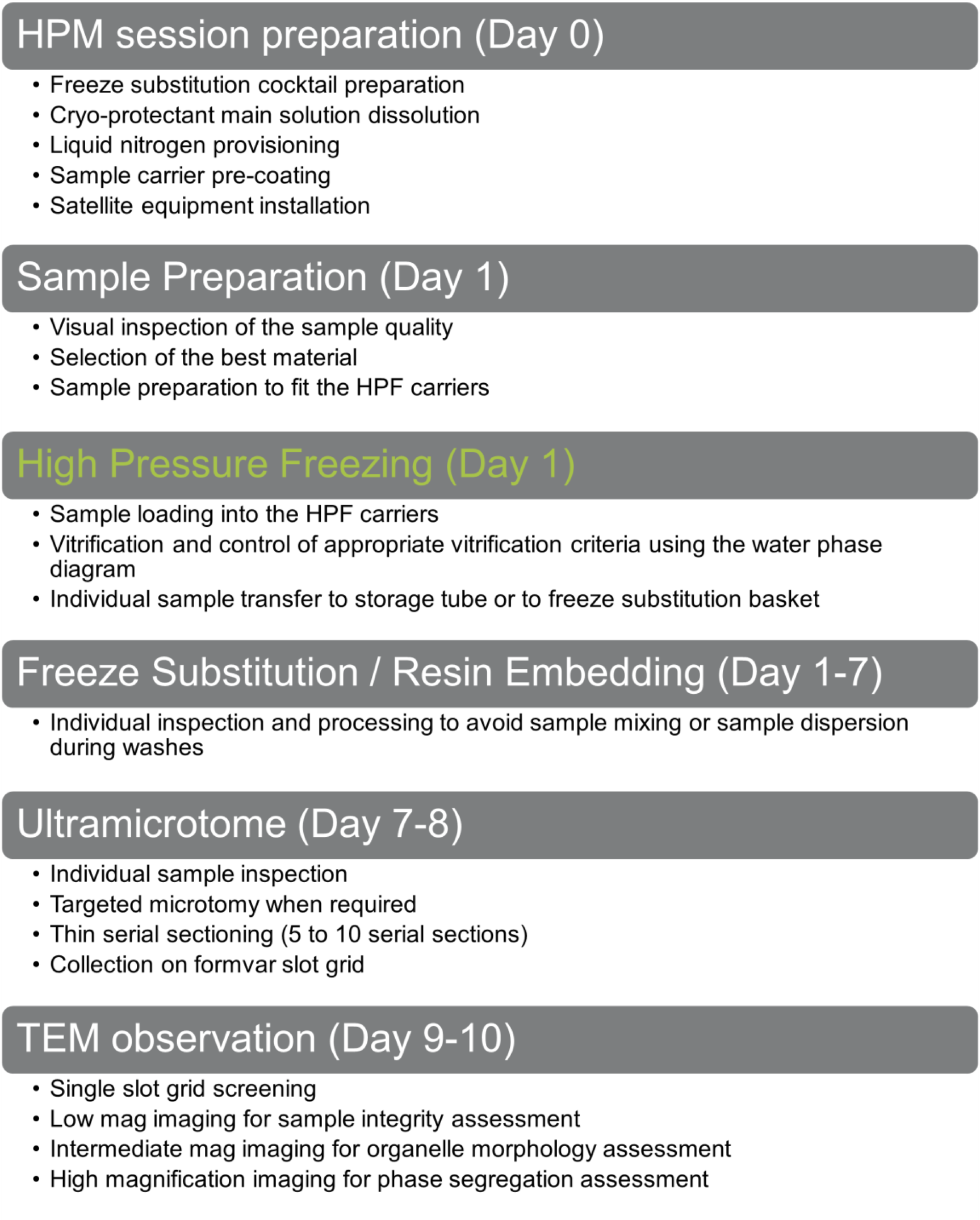
Global Workflow to assess sample vitrification versatility of our HPM Live µ over 9 different samples.

During vitrification sessions, meticulous attention was given to each sample’s journey across the water phase diagram, serving as a reliable indicator of HPF efficacy. Any samples showing signs of inadequate vitrification were promptly discarded and repeated, given the importance of this initial step in the process. We favor the water phase diagram display over conventional temperature and pressure crossing points, as it provides more informative insights and avoids potential inaccuracies due to uncalibrated scales.

Following this rationale, we conducted at least two independent HPF sessions with valid HPF curves for each sample type. In total, we present data from 25 independent experiments covering nine distinct sample types (including cell pellets, tissues, and small model organisms), showcasing the capability of our HPM Live µ to effectively vitrify a wide range of biological materials within the physical depth limitation of 200µm.

### Vitrification of human cell pellet

Three SHSY-5Y cell pellets (Samples A, B, and C) were prepared and subjected to high pressure freezing (HPF). The resulting HPF curves met the criteria for further processing for electron microscopy (Figure 2, line 1). Overview examination of the ultrastructure revealed well-preserved samples, without major ice nucleation causing large segregation (Figure 2, line 2). Observation of the culture medium mixed with PVP (sample A and B) displays a globally poor preservation, while the culture medium mixed with Dextran (sample C) shows no sign of ice nucleation (Figure 2, line 2). Sample B does not present a significant contrast increase despite higher concentration of osmium tetroxide during freeze substitution (Table 1). Sample C displays a lower contrast compared to sample A (same FS conditions). At intermediate magnification, organelles such as the Golgi apparatus, endosomes and mitochondria display regular morphology in all 3 samples (Figure 2, line 3). At higher magnification, continuous and sharp membranes observed across all samples and no segregation pattern appears in the cytoplasm (Figure 2, line 4).

**Figure 2:**
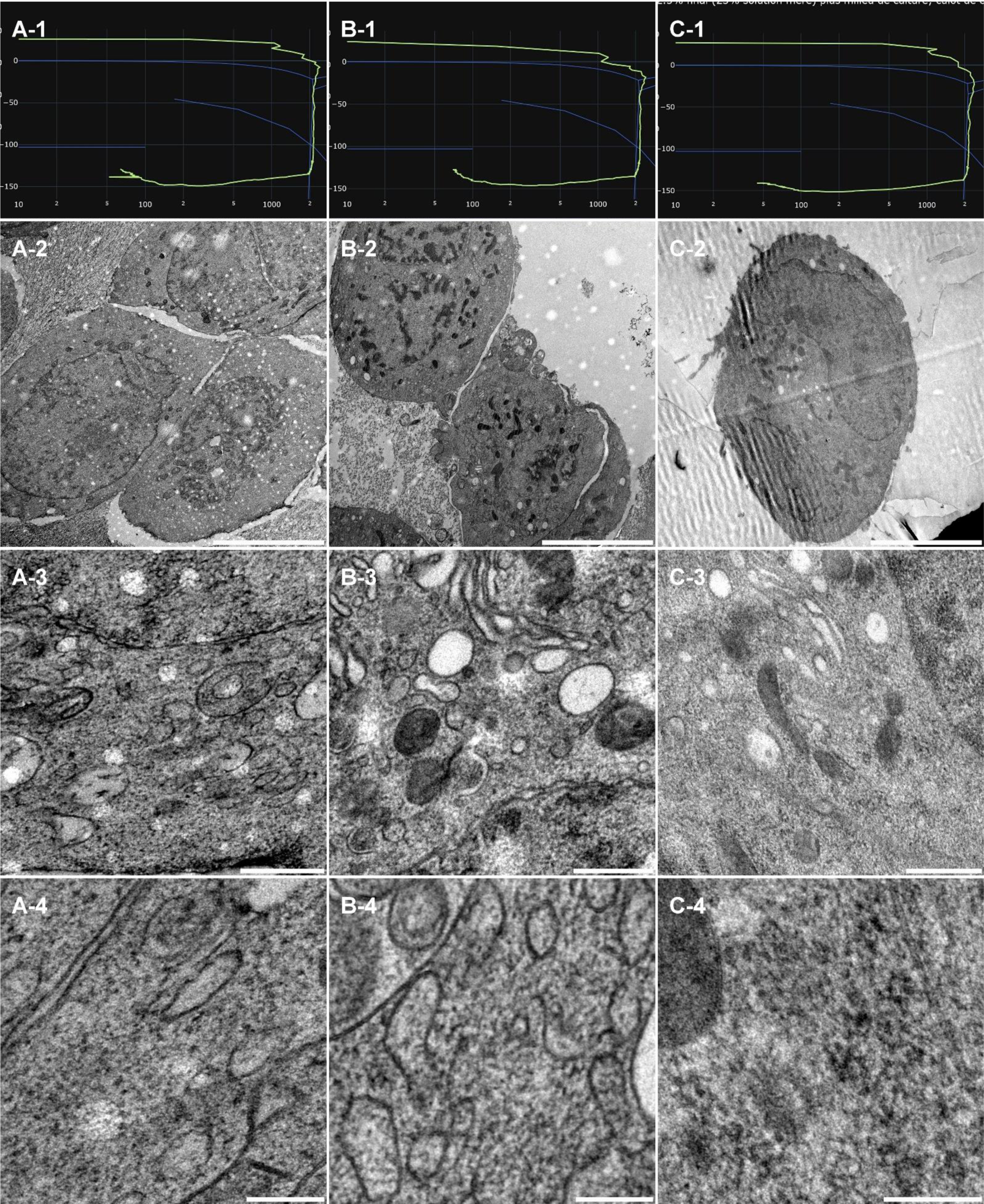
morphological observation of 3 samples (Column A, B and C) of human cell pellet, concentrated by centrifugation and resuspended in cryo-protectants. Line 1 present each individual HPF curve. Line 2 presents an overview of the samples, scale bar 5µm. Line 3 presents general organelle preservation and distribution, scale bar 0.5µm. Line 4 presents a higher magnification of a detailed view with endosomes (A, B), endoplasmic reticulum (B), mitochondria (A, C), cytoskeleton(C) and cytoplasm (A, B, C), scale bar 0.2µm.

**Figure 3:**
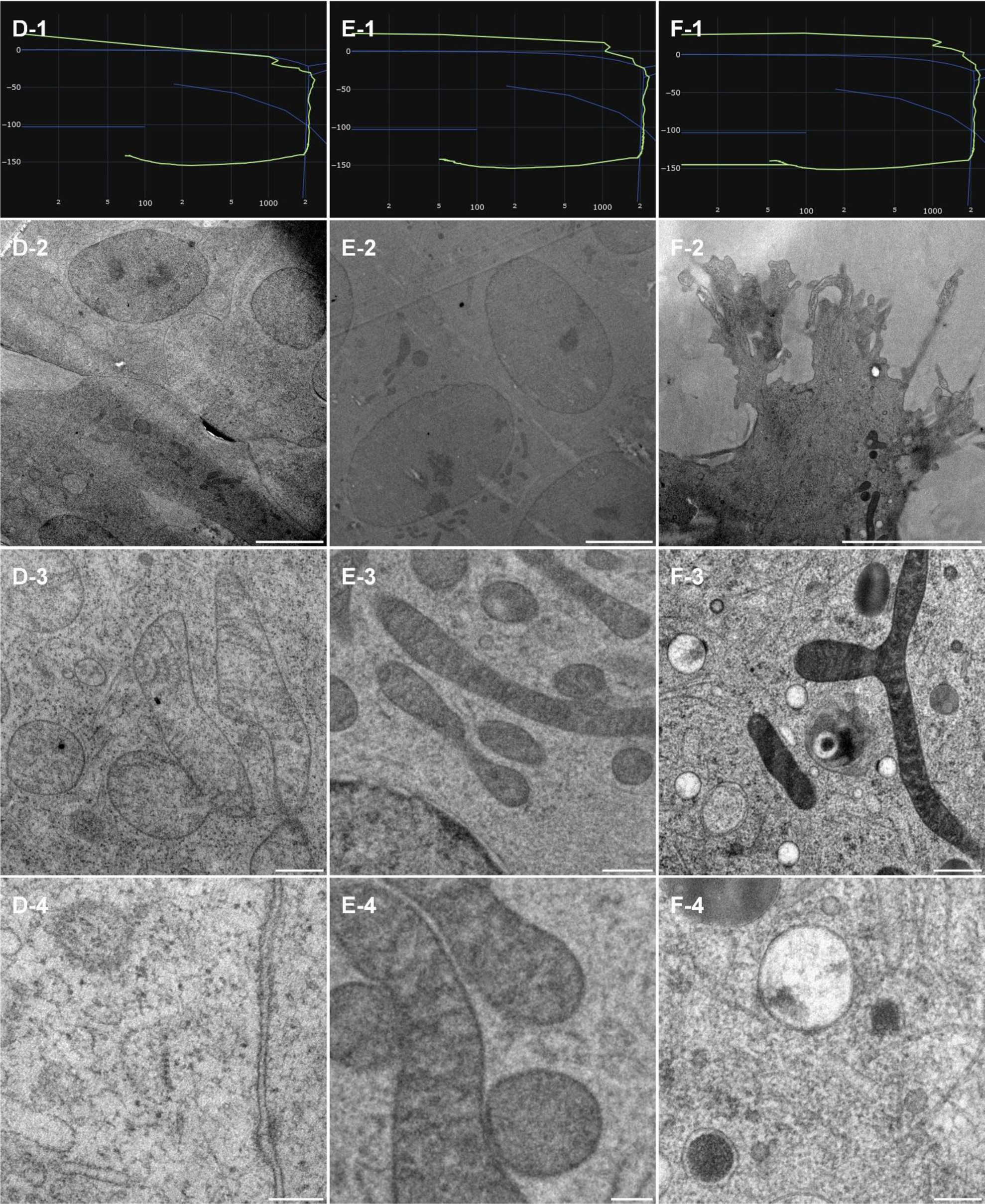
cell monolayer cultured onto CryoCapsule. Line 1 present each individual HPF curve. Line 2 presents an overview of the samples, scale bar 5 µm. Line 3 presents general organelle preservation and distribution, scale bar 0.5 µm. Line 4 presents a higher magnification of a detailed view with scale bar 0.2µm.

### Vitrification of cell monolayer on CryoCapsule

SHSY-5Y cells were cultured onto CryoCapsule (sample D, E and F), exposed to 8% dextran as a cryo-protectant and high pressure frozen. The resulting HPF curves of sample E and F met the criteria for further processing for electron microscopy. Sample D, despite a potentially bad HPF curve entering the solidification area 7ms before the critical 2000bars are achieved, was further processed out of curiosity. Overview examination of the cell culture revealed homogenously well-preserved samples. No crystallization between cells across the sample was observed. The culture medium mixed with dextran displays no evidence of ice nucleation and the nucleus are all nicely preserved. All freeze substitution were comparable, containing osmium tetroxide, and the contrast at the membranes is clearly delineated in all conditions. It is noticeable that mitochondria appear white (D-3) or black (E-3, F-3), and the overview picture present this variability in adjacent cells (D2, E2). These two CryoCapsules were freeze substituted on the same day. Sample F displays a stronger contrast at the mitochondria and was substituted on another day. Our apparatus for freeze substitution being manual, we suspect a time difference between reaching 4°C and the first acetone rinses.

**Figure 4:**
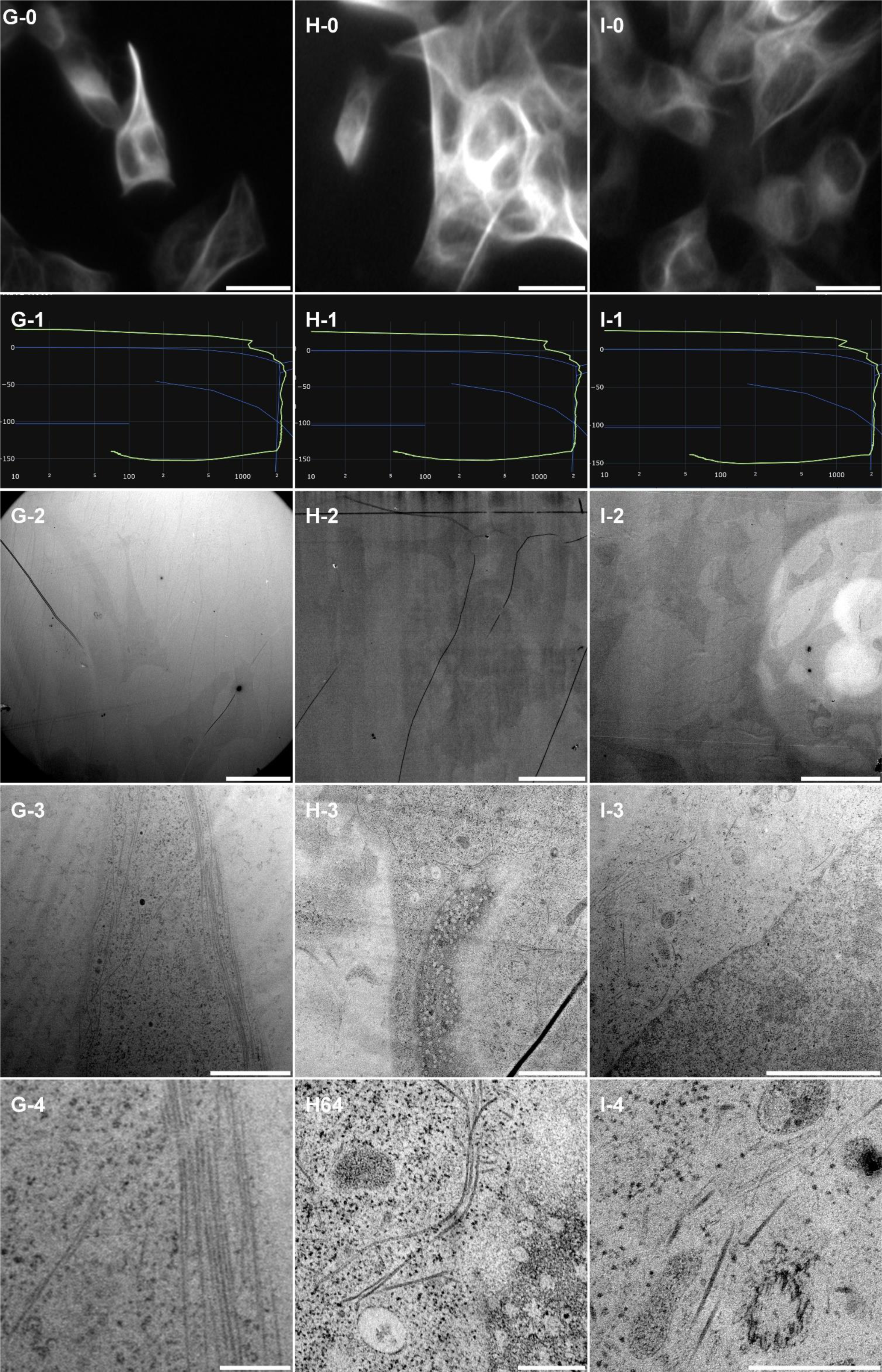
correlative light and electron microscopy analysis of cell monolayer cultured onto CryoCapsule. Line 0 present each ROI observed live on the HPM Live µ with a 20x magnification prior to HPF, scale bars 20µm. Line 1 present each individual HPF curve. Line 2 presents an overview of the samples, scale bar 20µm. Line 3 presents general organelle preservation and distribution, scale bar 2µm. Line 4 presents a higher magnification of microtubules, endosomes (G, H, I), mitochondria (H, I), nuclear pore complexes top view (H) and microtubule organizing center (I), scale bar 0.5µm.

### Vitrification of cell monolayer on CryoCapsule for CLEM analysis

SHSY-5Y cells were cultured onto CryoCapsule (sample G, H and I) to conduct live-CLEM experiments. To preserve sample preparation as close as possible to native, we did not add cryo-protectants to the sample prior to vitrification. The resulting HPF curves met the criteria for further processing for electron microscopy. In this experiment, we avoided osmium tetroxide in the freeze-substitution cocktail to avoid turning the whole sample black and facilitate sample retrieval at the ultramicrotome. As a consequence, membranes are not visible and only proteins inserted into them reveals their presence through the uranyl acetate stain. Cytoskeleton and proteins however are clearly observable. No discontinuity in cytoskeleton nor protein alignment along membrane is observed, highlighting a proper ultrastructure preservation of the sample.

From the CLEM perspective, we could locate all the cells observed live, back in the electron microscope, demonstrating the high reliability and reproducibility of the CryoCapsule for live-CLEM approached on cell monolayers.

### Vitrification of mouse brain biopsy

Three punches from perfused brain slices (Samples J, K, and L) were randomly selected for further vitrification. The resulting HPF curves of sample J and K met the criteria for further processing to electron microscopy (figure 5, line 1). Sample L, despite a potentially bad HPF curve entering the solidification area 7ms before the critical 2000bars are achieved, was further processed out of curiosity. Overview examination of the ultrastructure revealed homogeneous preservation across all 3 tissues with regular spacing between the myelinated neurons (Figure 5, Line 2). At intermediate magnification, organelles display a smooth and regular morphology (Figure 5, Line 3). It is interesting to note that the myelinated neurons do not appear densely packed as usually observed in volume EM dataset prepared by the OTO protocol[23]. This results from the gentler dehydration process obtained using freeze-substitution[24]. At higher magnification, individual myelin sheaths, dense vesicular regions, and mitochondrial cristae with double leaflet membranes are clearly discernible (Figure 5, Line 4).

**Figure 5:**
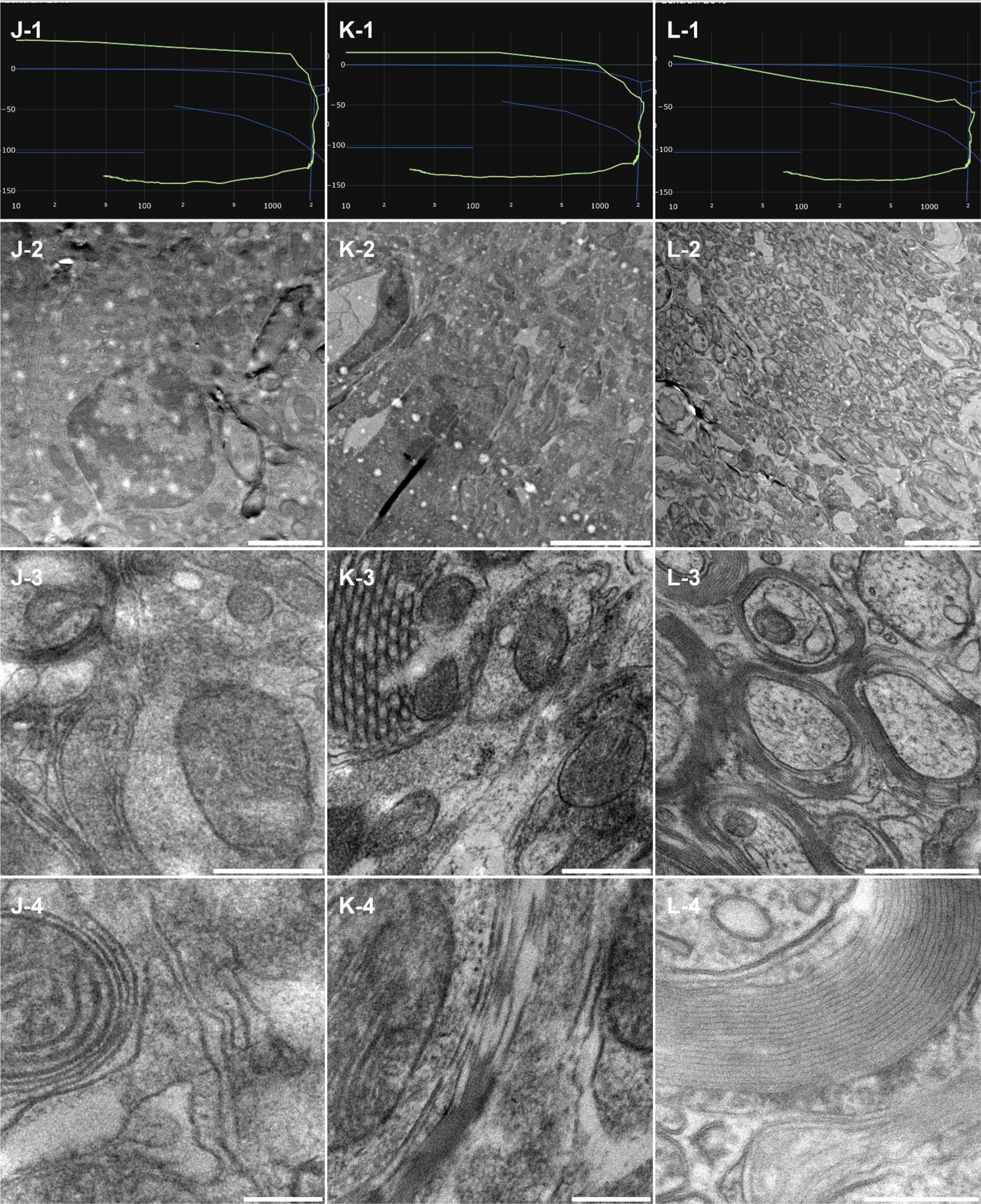
morphological observation of 3 samples (Column D, E and F) of mouse brain biopsies, sliced using an oscillating knife microtome. Line 1 present each individual HPF curve. Line 2 presents an overview of the samples, scale bar 5µm. Line 3 presents general organelle preservation and distribution, scale bar 0.5µm. Line 4 presents a higher magnification of a detailed view with membranes, myelin sheath, neuronal vesicles and cytoplasm, scale bar 0.2µm.

### Vitrification of mouse liver biopsy

Liver slices of 100µm thick were randomly selected and punched into three samples (M, N, and O). The resulting HPF curves met the criteria for further processing to electron microscopy (figure 6 line 1). Overview assessment of the ultrastructure revealed homogenous preservation across all tissues (Figure 6, Line 2). Sample M’s HPF curve was at the edge of the crystallization line, but remained in the liquid phase until proper pressure was attained. At all magnifications, sample M displays comparable morphological preservation to the other samples. Thereby confirming that the HPF parameters were met to achieve good ultrastructure preservation without ice nucleation.

**Figure 6:**
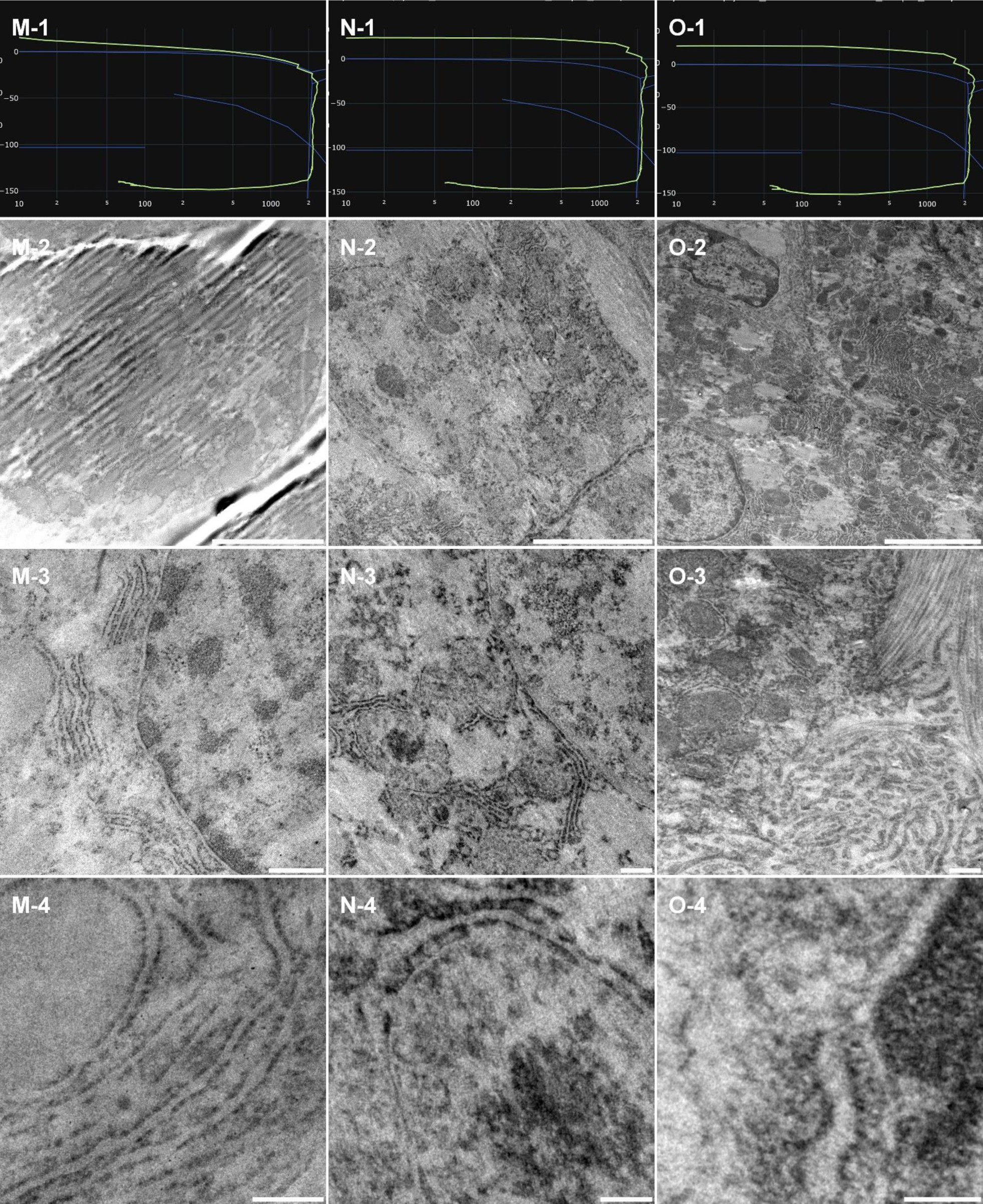
Morphological observation of 3 samples (Column M, N and O) of mouse liver biopsy, sliced using an oscillating knife microtome. Line 1 present each individual HPF curve. Line 2 presents an overview of the samples, scale bar 5µm. Line 3 presents well preserved organelles, scale bar 0.5µm. Line 4 presents a higher magnification of a detailed view of membranes from endoplasmic reticulum (M), glycogen compartment (N) and nuclear membrane (O), scale bar 0.2µm.

Sample M and N were freeze substituted without osmium tetroxide. As a result, the membranes are less prominently marked or appear lighter at higher magnification (Figure 6, line 4). Sample O was exposed to osmium tetroxide during freeze-substitution, resulting in darkly stained membranes.

In all three samples at intermediate magnification, distinct morphologies were evident, with no clear ice segregation patterns (Figure 6, Line 3). At higher magnification, sharp and continuous structures were observed across all samples (Figure 6, Line 4).

### Vitrification of C. elegans

Three *C. elegans* samples (P, Q and R) were concentrated (see table 1) and deposited in 200µm deep carriers. The resulting HPF curves met the criteria for further processing to electron microscopy (figure 7 line 1).

**Figure 7:**
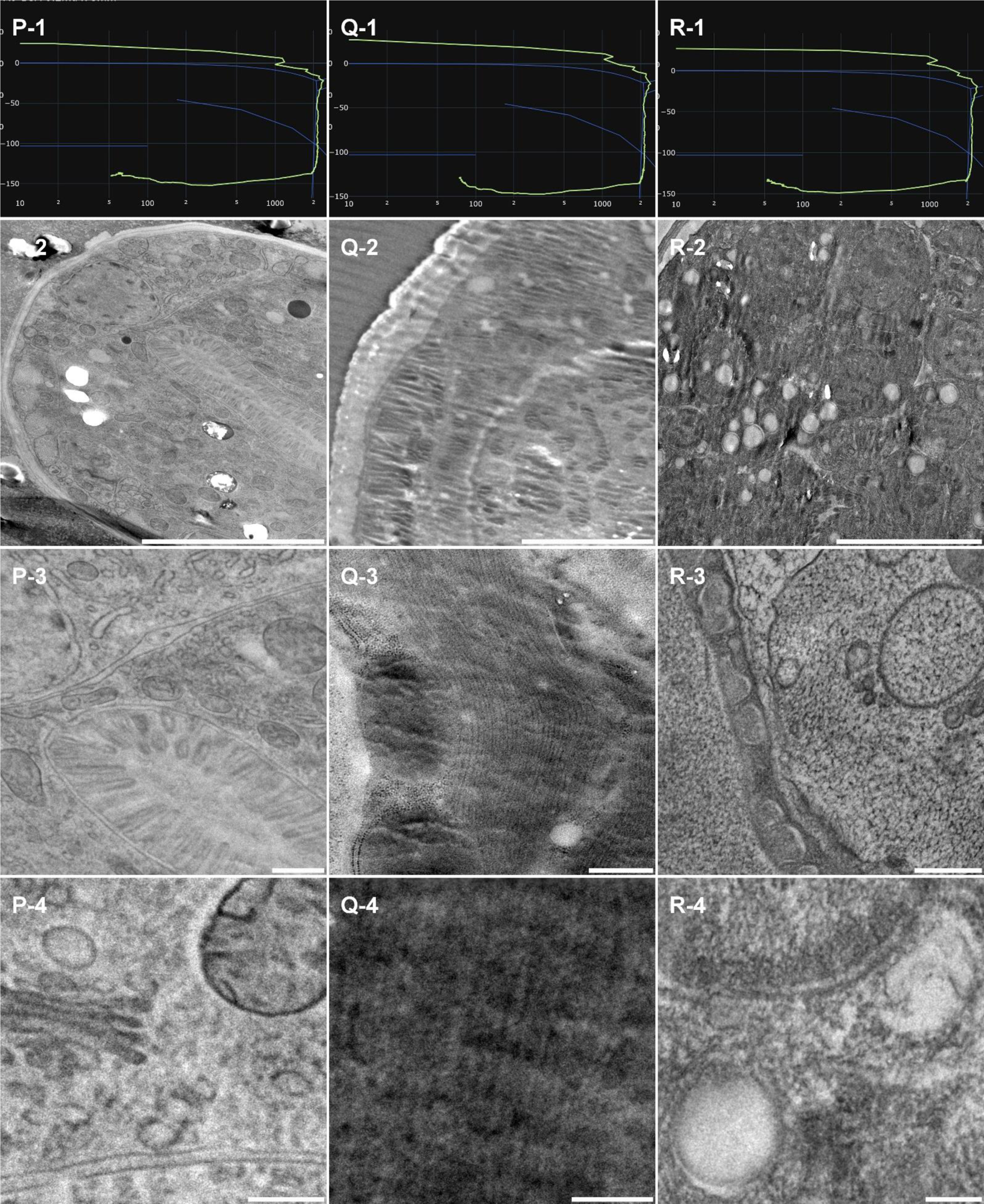
morphological observation of 3 samples (Column M, N and O) of *Caenorhabditis elegans*. Line 1 present each individual HPF curve. Line 2 presents an overview of the samples, scale bar 5µm. Line 3 presents well preserved organelles, scale bar 0.5µm. Line 4 presents a higher magnification of a detailed view of membranes from Golgi, mitochondria (P), muscle fibers (Q) endosomes and nuclear membrane (R), scale bar 0.2µm.

**Figure 8:**
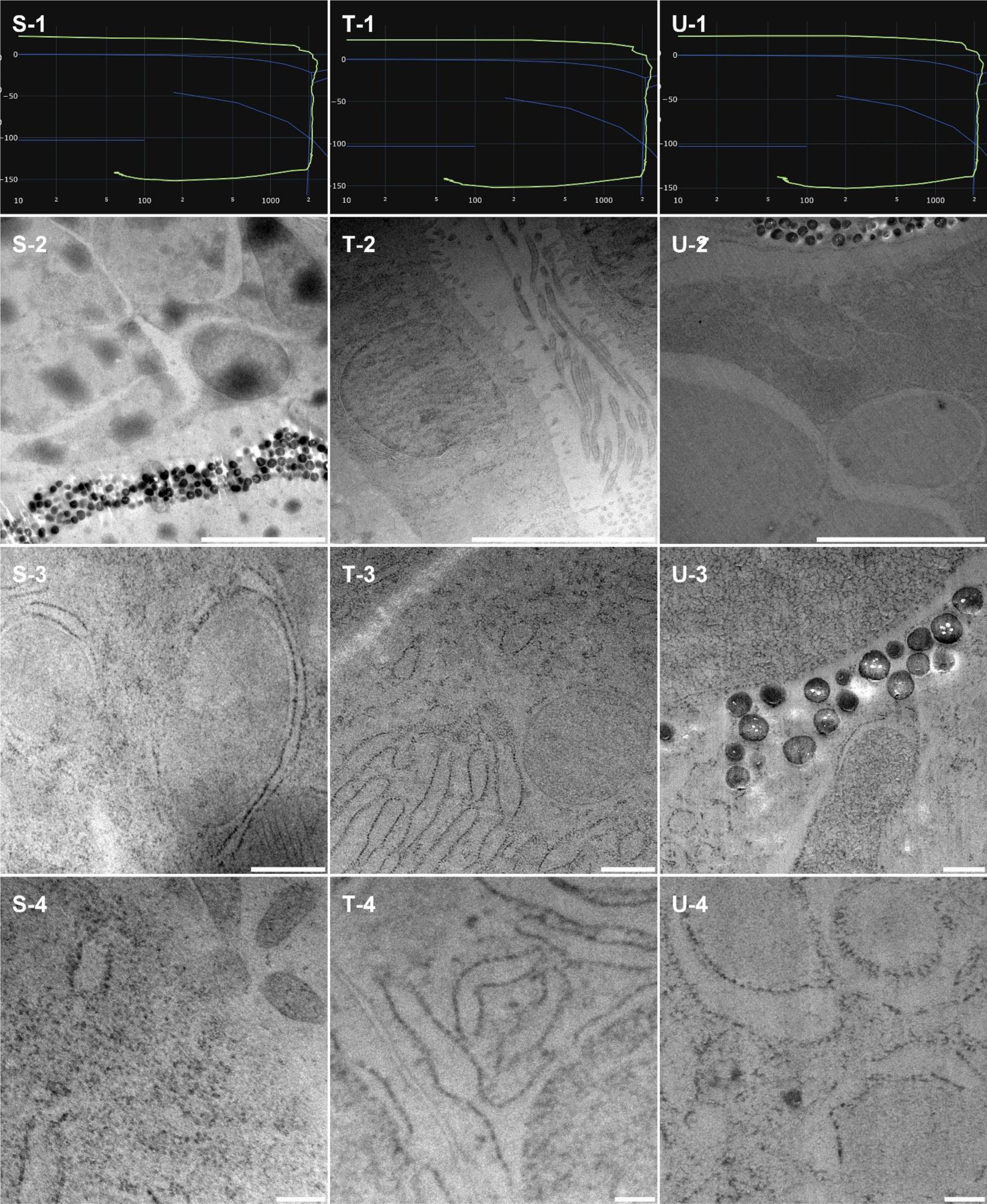
morphological observation of 3 samples (Column M, N and O) of *Danio rerio*, Zebrafish. Line 1 present each individual HPF curve. Line 2 presents an overview of the samples, scale bar 5µm. Line 3 presents well preserved organelles, scale bar 0.5µm. Line 4 presents a higher magnification of a detailed view of membranes from endoplasmic reticulum (S, T, U) and nuclear envelope (T), scale bar 0.2µm.

Sample P was freeze substituted in presence of 1% osmium tetroxide with the usual temperature ramp (2°C.hr).

Sample Q was freeze substituted in presence of 1% osmium tetroxide but underwent a faster temperature ramp (4°C/hr)

Sample R was freeze substituted in presence of 2.5% osmium tetroxide with the usual temperature ramp (2°C/hr)

Overview assessment of the ultrastructure revealed homogenous preservation across all tissues (Figure 7, Line 2).

Contrast in sample Q seems milder than contrast of P and R. Interestingly, sample R does not seem to display a more prominent contrast than sample P despite the higher amount of osmium tetroxide. This observation is redundant with the cell pellet contrast difference.

At intermediate magnification (figure 7, line 3), the ultrastructure appear homogenous and well preserved. At higher magnification (figure 7, Line 4), we observe sharp continuous membranes and nicely preserved ultrastructure,

### Vitrification of zebrafish

In our early experiments, we used osmium tetroxide to get comparable freeze-substitution parameters to the other experiment. However, after freeze substitution, the entire pellet turned black, and we could not distinguish the location of the zebrafish larvae in the EPON bloc. Random sectioning being too time consuming and too hazardous, we decided to remove osmium tetroxide in the following freeze substitution cocktails. As a direct consequence, the contrasts were globally low in the three samples presented here (S, T and U). Furthermore, sample U was treated with lower concentration of UA (Table 1).

From a lower perspective, the sample had more heterogenous preservation quality. The sample is extremely large and must be gently squeezed between the aluminium planchettes. Aside from the physical stress, the carrier must be filled with a significant volume of Dextran 20% to reduce the water content surrounding the sample. It may affect the physiology of the sample prior to vitrification and a special attention must be taken by the experimentalist as of the scientific question to ask. A pre-fixation might be wise to consider.

Yet, we observed several well-preserved areas in all three samples near the eye, or around the muscles. Extensive investigation across the entire fish volume to estimate the best-preserved parts is out of reach within our current facility.

At intermediate magnification (figure 7 line 3), we observe organelles with smooth and homogenous distribution.

At higher magnification (figure 7, line 4) we observe smooth continuous membranes and nicely preserved ultrastructure.

### Vitrification of A. thaliana root

From the global view, the ultrastructure of the two samples (V and W) is well preserved and homogenous. Sample V displays the typical plant root structure with organized cell arrangement and large vacuoles (Figure 9, V-2). For sample W, we could not find the tissue structure like V, but inner cellular ultrastructure was well preserved (Figure 9, W-2). at intermediate magnification (Figure 9, line 3) we observed well organized and preserved organelles. At higher magnification (figure 9, line 4) we observed smooth continuous membranes.

**Figure 9:**
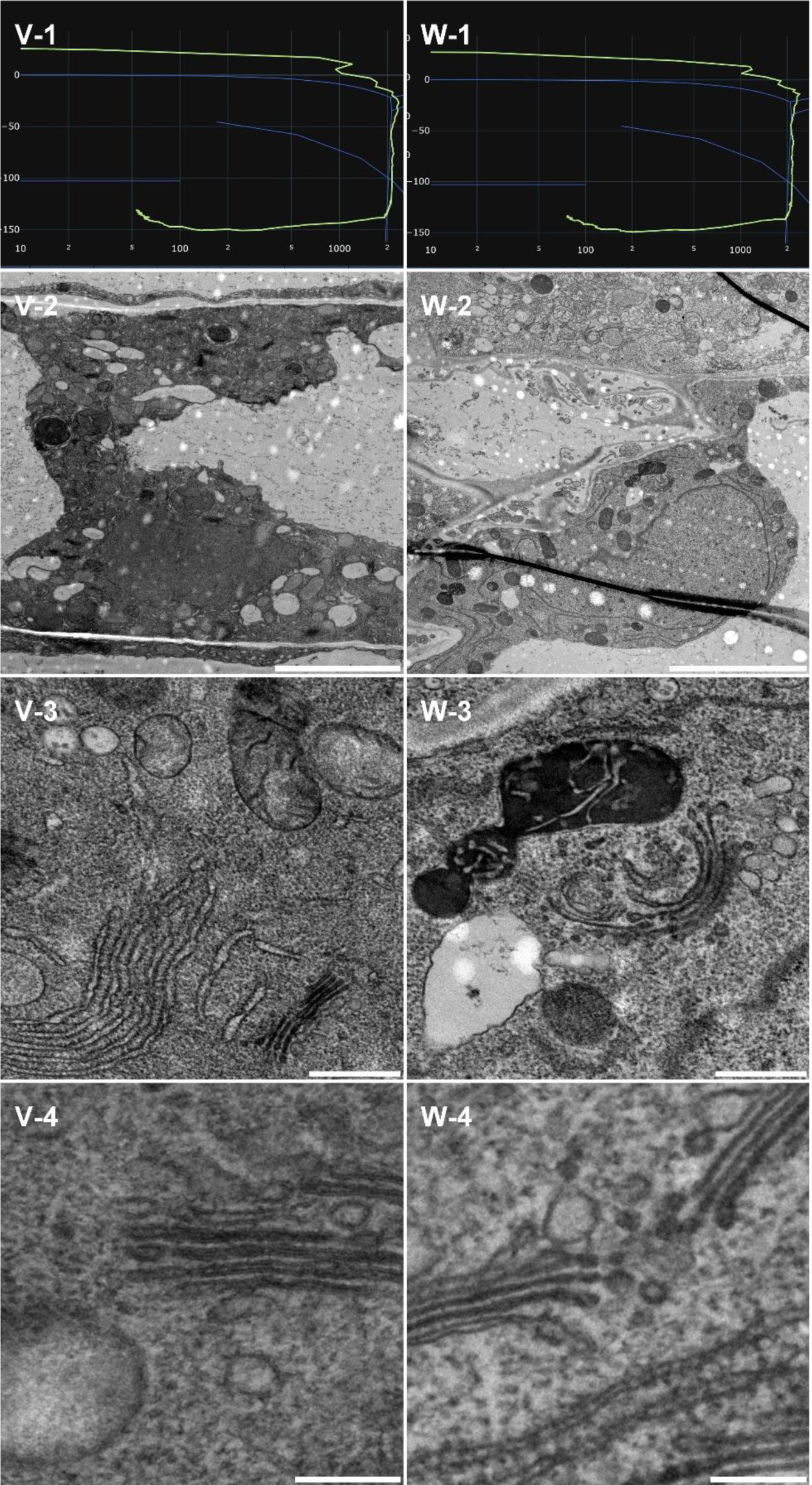
morphological observation of 2 samples (Column M, N and O) of Arabidopsis thaliana root. Line 1 present each individual HPF curve. Line 2 presents an overview of the samples, scale bar 5µm. Line 3 presents well preserved organelles, scale bar 0.5µm. Line 4 presents a higher magnification of a detailed view of membranes from golgi apparatus and endosomes (V, W), scale bar 0.2µm.

### Vitrification of A. thaliana seedling

Plant leaves are notably challenging to vitrify. The presence of large air vacuoles prevents the proper building of the pressure up to 2100 bars which is necessary to achieve fast glass transition and limit ice nucleation. In this figure, we present two samples (X and Y) that were well preserved. We observe from the large perspective (Figure 10, line 2), some membrane ruptures and sample fractures. This likely caused by mechanical rupture around the vacuoles during vitrification.

**Figure 10:**
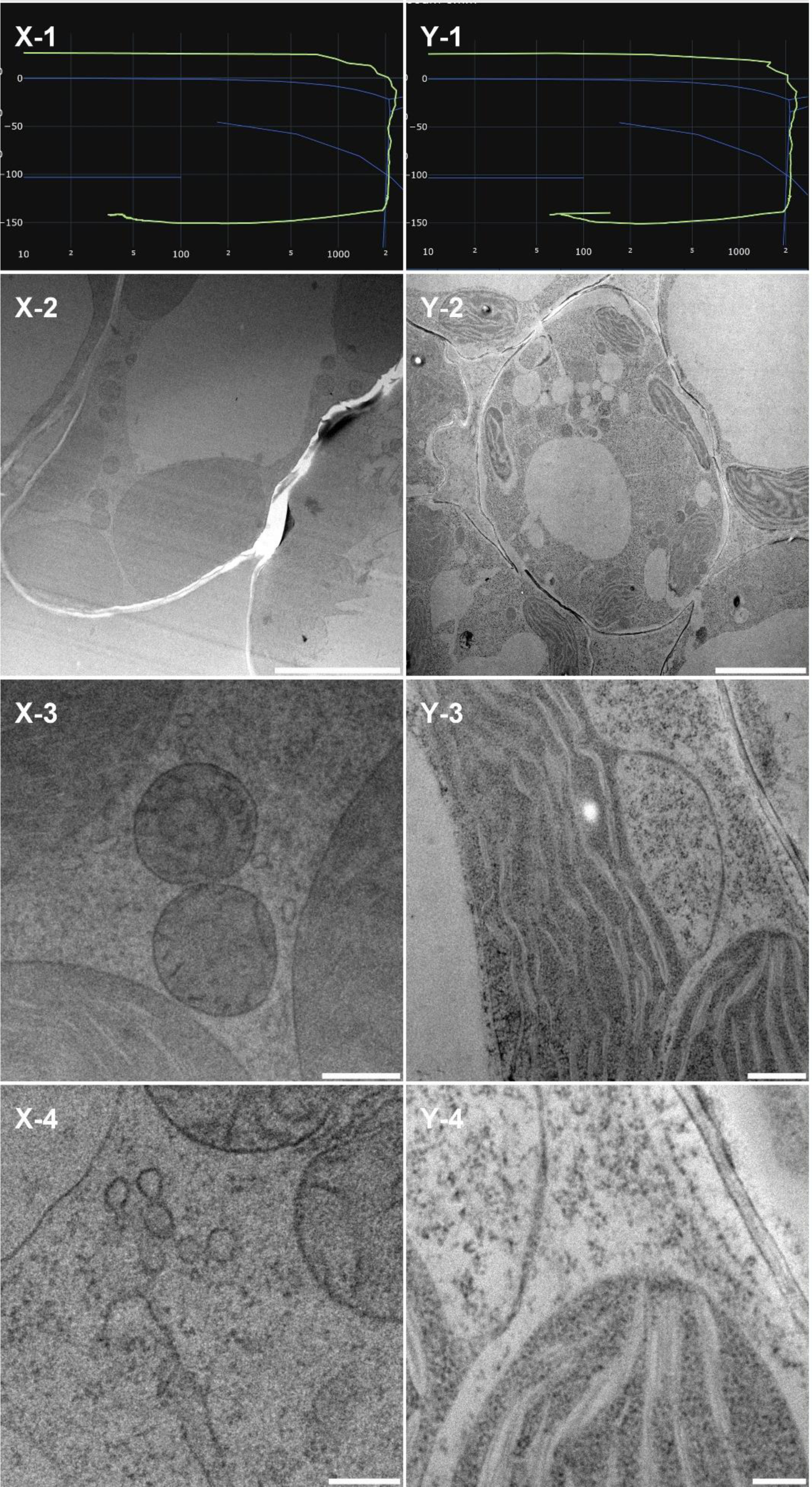
morphological observation of 2 samples (Column M, N and O) of Arabidopsis thaliana seedling. Line 1 present each individual HPF curve. Line 2 presents an overview of the samples, scale bar 5µm. Line 3 presents well preserved organelles, scale bar 0.5µm. Line 4 presents a higher magnification of a detailed view of mitochondria, endosome (X) and thylakoid, cytoplasm (Y), scale bar 0.2µm.

However, closer inspection (figure 10, line 3) displays nicely preserved organelles typical of this plant tissue, the chloroplast (Figure 10, Y-3). At higher magnifications, trafficking vesicles and other membrane structures were very smoot and distinguishable (Figure 10, D and H).

## Discussion

In the realm of biological specimen preservation, high pressure freezing (HPF) stands out as the exclusive method capable of vitrifying specimens ranging from 5µm to 200µm in thickness. Since its introduction in laboratories in the late 1980s, HPF has recently garnered renewed interest from the scientific community, particularly with the emergence of cryo-electron microscopy and volume electron microscopy spurred by the resolution revolution.

The HPM Live µ discussed in this article represents a significant advancement from the established HPM010 by BalTec, renowned as the pioneering and most dependable apparatus in the market. To validate the reliability of this newer iteration, we conducted extensive testing on a diverse array of biological specimens, including isolated cells, tissues, and small model organisms. Our evaluation encompassed sample preservation quality at varying scales, from multiple cells to individual organelles, employing freeze substitution and room temperature imaging techniques.

By leveraging freeze substitution and room temperature imaging, we were able to comprehensively assess sample preservation homogeneity across different locations, up to millimeters apart on a formvar slot grid. High-magnification observations enabled the identification of microcrystalline ice-induced damages such as membrane rupture or protein segregation.

This sample preparation and imaging strategy facilitated the exploration of a wide range of samples, irrespective of prior expertise. By establishing comparable assessment criteria and employing widely used protocols, we aim to provide facility laboratories with a foundational framework for handling novel samples. Our detailed protocol serves as a reference point, enabling researchers to gauge the adequacy of their own sample preparations. Nonetheless, we acknowledge that sample preparation may require further refinement, with post-processing techniques like freeze-substitution tailored to specific research objectives, such as membrane analysis or cytoskeletal studies, drawing upon existing literature knowledge.

This collaborative endeavor involved two distinct scientific profiles: Chie Kodera, PhD, a novice in electron microscopy who acquired experimental skills over the course of a year, and Xavier Heiligenstein, PhD, a seasoned expert in high pressure freezing and correlative microscopy techniques, who provided training and project supervision. Together, we established rigorous procedures to establish a novel HPF service at the INSERM Unit U1195 in Le Kremlin-Bicêtre, France.

Our two-year project underscored the benefits of collaborative sample preparation, emphasizing the advantages of having one individual dedicated to sample preparation and loading while another focuses on collection and storage. When specialized expertise is required for sample preparation, end users must oversee this process before transferring samples to HPF carriers.

Given the noisy and delicate nature of HPF apparatus and the typically crowded laboratory environments, we advocate for thorough preparation the day before an experiment, including the preparation of sample carriers, cryoprotectant master solutions, freeze-substitution cocktails, and apparatus precooling. This approach ensures that the experiment day can be dedicated solely to sample handling and data collection.

Following our established procedures and experimental protocols, we conducted HPF on a diverse range of biological specimens using the HPM Live µ apparatus. We refined various preparation protocols and present the most conclusive ones here.

Thinning samples often necessitated real-time adjustments by the experimentalist. We recommend practicing on dummy samples one or two days prior to the actual experiment, enabling the identification of necessary materials and the development of effective techniques.

While cryo-protection is ideally avoided, it can be beneficial when exploring novel or challenging samples. On pre-fixed samples, cryo-protection causes minimal osmotic shock effects, and helps preserving sample ultrastructure homogeneity.

Our selection of EPON embedding resin for all samples was primarily based on its widespread use and established reputation for embedding various biological specimens with consistent quality.

However, it is important to note that EPON resin tends to reduce the contrast generated by the heavy metal salts present in the freeze substitution cocktail, often necessitating post-staining procedures for enhanced visualization. In our study, we deliberately avoided post-staining to minimize variability and showcase the true effectiveness of each sample preparation protocol.

An evident consequence of this choice was observed in samples that did not undergo osmium tetroxide exposure during freeze substitution, resulting in notably weak contrast, especially in membrane structures, which uranyl acetate alone could not adequately compensate for. Osmium tetroxide, while effective for contrast enhancement, poses significant challenges in terms of health and safety regulations, as well as disposal requirements. Additionally, it can cause the entire sample pellet to turn black, as seen in our CLEM on CryoCapsule and zebrafish experiment, hindering the identification of specific samples for targeted microtomy.

Addressing this technical challenge presents complexities, as alternatives such as soft X-ray apparatus, while capable of mitigating osmium tetroxide-related issues, are not universally available and entail substantial costs, making them impractical for routine screening protocols like the one presented in our study. To circumvent these challenges, we propose the consideration of more electron-transparent resins, such as HM20 Lowicryl® [25](EMS, USA), or contrast-enhancing electro-conductive resins like R221® [26](CryoCapCell, France), which could offer improved visibility of cellular structures without compromising sample integrity.

During the two-year duration of this study, we thoroughly analyzed various aspects of the high-pressure freezing (HPF) protocol, including pre-fixation, cryoprotection, HPF curve evaluation, freeze substitution cocktail impact, and the influence of resin on contrast (though not presented in this manuscript). While meeting the expected vitrification criteria, namely reaching 2000 bars before cooling and achieving a temperature decrease rate of over 2000K/s, is crucial, we deliberately examined samples that did not meet these criteria out of scientific curiosity, as exemplified by samples D and L. To our surprise, these samples still exhibited high-quality ultrastructure under scrutiny with an electron microscope.

Several interpretations can be proposed to account for these irregularities. Firstly, the positioning of the thermocouple sensor, located at the top of the HPF chamber, and the pressure sensor, situated at the bottom, may introduce a slight delay in the readout. The direct path of the liquid nitrogen jet to the sample, followed by its indirect path to the sensors, could result in asynchronous temperature and pressure measurements, potentially affecting the observed vitrification outcome.

Secondly, the use of ethanol in the HPF chamber prior to the release of liquid nitrogen, known to delay sample cooling while pressure rises to the desired 2000 bars, may play a role. Even in small amounts, ethanol presence could synchronize temperature and pressure at the sample site independently of the built-in sensors, possibly influencing the vitrification process.

Lastly, the patented pressurization technology of the HPM Live µ, enabling rapid pressure rise to 2000 bars in less than 10ms, compared to its predecessor, the HPM010 [27], [28], which required ethanol delay with pressure rise taking up to 25ms, could be a contributing factor. Deciphering between these interpretations presents our next challenge in optimizing sample vitrification by high-pressure freezing. Further investigation and experimentation are warranted to elucidate the underlying mechanisms and refine our understanding of the HPF process.

## Conclusion and outlook

In summary, our study demonstrates the preservation of high-quality ultrastructure across a broad range of samples, meeting the requirements of most facilities. The sample preparation parameters proposed here are likely applicable to all high-pressure freezing apparatuses. By ensuring that samples undergo the appropriate journey across the water phase diagram, researchers can use this manuscript as a guide to establish internal HPF protocols.

## Acknowledgement

CK and XH conceived the project, ran the experiment, and wrote the manuscript, YB, FE, XH and JH conceived the HPM Live µ, MB supervised the project.

We thank Yvette Akwa (INSERM UMR 1195) and Kevin Guillemeau (INSERM UMR 1195) for neuroblastoma cells, Christian Specht (INSERM UMR 1195) for mouse brain tissue, Thibaud Sefiane (INSERM UMR 1176), Marie Clavel (INSERM UMR 1176), and Peter Lenting (INSERM UMR 1176) for mouse liver tissue. Ophélie Nicolle (Université de Rennes) and Grégoire Michaux (Université de Rennes) for *C. elegans*, Cindy Degerny (INSERM UMR 1195) and Marcel Tawk (INSERM UMR 1195) for Zebrafish, Claire Boulogne (Plateforme Imagerie-Gif Institut de Biologie Intégrative de la Cellule, UMR 9198) for *A. thaliana*, RMC Boeckeler Instrument, and the Francis Crick institute for the kind lending of the RMC FS8500 freeze substitution apparatus. We acknowledge the ImagoSeine core facility of the Institut Jacques Monod, member of the France BioImaging infrastructure (ANR-10-INBS-04) and GIS-IBiSA” for the use of TEM.

## Notes

### Competing Interest Statement

The authors are the manufacturers and distributors of the HPM Live μ, used in this manuscript.

